# A framework for Polinton-like virus diversity reveals links to multiple viral classes and Nucleocytoviricota

**DOI:** 10.64898/2026.06.19.733378

**Authors:** Christopher Bellas, Ruben Sommaruga

**Affiliations:** Department of Ecology, Universität Innsbruck, 6020 Innsbruck, Austria; Department of Ecology, University of Innsbruck, Technikerstrasse 25, A-6020 Innsbruck, Austria

**Keywords:** Polinton, Maverick, Virophage, NCLDV, Nucleocytoviricota

## Abstract

Polinton-like viruses (PLVs) are among the most abundant eukaryotic DNA viruses in aquatic environments. Despite their extensive diversity, broad host range and variable gene content, they are commonly treated as a single group, which obscures their evolutionary relationships and complicates their classification.

Through analysing thousands of viral genomes from aquatic ecosystems and public metagenomic datasets, we clarify the evolutionary structure encompassed by the term PLV. Using sensitive profile Hidden Markov Model (HMM) comparisons, phylogenies of conserved capsid morphogenetic genes and gene content analysis, we show that viruses referred to as PLVs are distributed across multiple deep lineages spanning at least three currently recognised viral classes. These include the Gosseviruses, aquatic viruses related to Maverick-Polintons in animal genomes. They also include a continuum of related viruses from 15 kb PLVs to the 45 kb Mriyaviruses and more broadly, to the Nucleocytoviricota, potentially representing extant relatives of giant viruses. Our findings suggest that PLVs do not fit neatly within existing taxonomic boundaries, reflecting a complex history of horizontal gene transfer and diversification of life strategies. To support future discovery, we provide a curated set of HMMs representing the known capsid diversity of PLVs, Maverick-Polintons, and virophages. This toolkit enables sensitive detection and identification of PLVs across metagenomic and eukaryotic genome datasets.

Our study provides an evolutionary framework for interpreting PLV diversity and a foundation for future refinement of their classification.

## Introduction

The discovery of small, eukaryote-infecting DNA viruses in the 15-45 kb size range has accelerated in recent years, offering new insights into virus evolution and eukaryotic genomics. However, there remains considerable variability in the literature regarding how to define many of these viruses and organize their taxonomic relationships. Initially, 15-20kb elements called Mavericks or Polintons (hereafter referred to as Maverick-Polintons), which encoded protein-primed DNA polymerase B (pPolB) and retroviral integrases (RVE), were discovered and catalogued as transposable elements in eukaryotic (mainly animal) genomes [1,2]. These were later reclassified as viruses having the genetic potential to construct a viral capsid and package their DNA through the possession of virus capsid and packaging genes (Major Capsid Protein - MCP, minor Capsid Protein - mCP, genome packaging A32-like ATPase, capsid maturation protease) [3]. Although no virions have been observed for Maverick-Polintons, they have been extensively described in vertebrates [4], with the remnants of ancient integrations even preserved in the Human genome [5–7]. Maverick-Polintons are also implicated as one of the long-theorised vectors for horizontal gene transfer in eukaryotes. In studies of nematodes, there is evidence for their horizontal transfer along with nearby host genes, suggesting encapsidated viruses move genes across species [8,9]. Some Maverick-Polintons are also referred to as ‘Adintoviruses’ [5], now consolidated by the ICTV under the Class *Polintoviricetes* [10]. Hence, despite different naming conventions between different research fields, the terms Mavericks, Polintons, Adintoviruses, and Class *Polintoviricetes* generally refer to the same viral elements.

Related to Maverick-Polintons are the Polinton-like viruses, initially discovered through metagenomic and genomic analysis [11,12] and later found to be highly diverse and abundant in aquatic ecosystems [13]. PLVs are similar in length to Maverick-Polintons, possessing a shared core viral morphogenic module (MCP, mCP and A32-like ATpase), however these core genes are highly divergent from the original Maverick-Polintons, with little detectable sequence similarity. The remaining PLV gene content is also highly variable, with pPolB genes often replaced by DNA Polymerase-Helicase fusion genes for replication, and RVE integrases replaced by Tyrosine recombinases for genomic integration. PLVs form several clusters by analysis of shared gene content and MCP phylogeny [13], suggesting PLVs as a whole represent multiple groups of viruses which await classification. PLVs are abundant in the genomes of protists, with tens to thousands of virus integrations in individual genomes [14–16], hence they appear to exist as both free virus particles and integrated virus forms. Only a handful of isolates have been described in detail in the literature, with many candidates awaiting investigation. Gezel (or PgVV), replicates only in the presence of a giant virus (*Nucleocytoviricota*) infecting the algae *Phaeocystis globosa*, hence has a virophage-like life strategy and is parasitic of a giant virus [16]. Similarly, at least one of three PLVs co-isolated with giant viruses of the algae *Chrysochromulina parva* is active against the giant virus [17]. The PLV *Tetraselmis striata* Virus was originally isolated as a nuclear replicating lytic virus, [18], however three closely related viruses Tsv-S2a, Tsv-S2b and Tsv-S3b are strictly dependant on a co-infecting giant virus for their replication, interestingly the latter two are reactivated endogenous elements from within the host genome [19]. Together this evidence opens up the possibility that many PLV groups represent viruses dependent on coinfecting giant viruses for their replication which have a viophage-like effect.

The discovery of Yaravirus, a lytic *Acanthamoeba* spp. infecting virus, has further expanded the diversity of small eukaryotic DNA viruses [20]. Yaravirus is the only isolated member of the Mriyaviruses [21], a group of 35-45kb viruses which include endogenous viruses detected in Chlorophyte genomes, referred to as NCLDV-dwarf-like viruses [16]. Mriyaviruses possess several giant virus (nucleocytoplasmic large DNA viruses - NCLDV) orthologous genes and NCLDV-like virus morphogenic modules, suggesting they are the closest known extant relatives of giant viruses [21], despite being in overlapping size ranges with the PLV. The distinction between Mriyaviruses and PLV is based on the presence of several core Mriyavirus genes, and the possession of a HUH endonuclease, indicative of rolling circle replication [21]. Hence, this separation from PLV, but similarity to giant viruses means Mriyaviruses are currently recognised as the smallest members of the *Nucleocytoviricota*.

Recent discoveries have further accelerated the discovery of Maverick-Polintons, PLVs and related elements. In a study of the Southern Ocean, abundant and diverse groups of PLVs have been detected and recognised as major components of marine viromes [22]; hundreds of Marick-Polintons and PLVs have been found integrated into the genomes of stony corals which show evidence of gene expression [23]; over a thousand PLVs have been detected in long-read assemblies of cryptophyte genomes [24], and PLVs have been found in the occlusion bodies of entomopoxvirus in insects, suggesting they are the first known parasites of poxviruses [25]. Clarifying the relationships between the PLVs is now essential to consolidate our understanding and begin to study the functional role of the many different virus groups.

*Polisuviricotina* is the recently defined supergroup (subphylum) of eukaryote-infecting DNA viruses that includes Maverick-Polintons (*Polintoviricetes*), Polinton-like viruses (PLVs), adenoviruses, and virophages (*Virophaviricetes*) [10]. In this framework, the PLVs are considered as a single group and hence, their internal diversity remains underexplored and their evolutionary relationships to other *Polisuviricotina* members are not fully resolved. Uncovering new PLV lineages and clarifying their relationships is essential for understanding their host range, potential for horizontal gene transfer, and diverse replication strategies. In this study, we systematically resolve the relationships between PLVs by integrating core gene phylogenies, gene content analysis and sensitive profile-based searches. In doing so, we reveal multiple deep viral lineages, some of which are more closely related to members of the *Nucleocytovirocota* than *Polisviricotina*. By unpacking the catch-all term ‘PLV,’ we establish an evolutionary framework and provide a toolkit to support future taxonomic efforts and refinement of the PLV structure.

## Results and discussion

In this analysis, we retrieved over 20,000 PLV genomes greater than 10kb in length from locally generated metagenomes from Austrian lakes and public sequence databases, greatly expanding the known diversity of PLV. By employing an iterative search strategy, we further detected over 70,000 MCP genes across all contig sizes, many of which defined new virus groups, allowing large-scale evolutionary relationships between the PLV to be established.

The diversity of double jelly-roll fold MCP genes encoded by PLVs is challenging to work with, MCPs from one group cannot be used to detect those from another using standard sequence similarity, or often even searching against single HMM profiles. To overcome this challenge, we performed sensitive all-vs-all HMM profile comparisons (HHsearch) of clustered MCP genes, producing a dissimilarity matrix of bitscores (Methods) to reveal distant relationships between PLV groups (Figure 1). This broad level comparison allowed the relationships of four highly diverse virus groups to be visualised: *Nucleocytoviricota* and related viruses, virophages (*Virophaviricetes*), Maverick-Polinton-related viruses (*Polintoviricetes*), and all remaining PLVs (*Aquintoviricetes*), which provided a framework for more in-depth phylogenetic analysis using ATPase genes. Viruses in this analysis, and those described previously in the literature as PLVs, fell into three of these groups using ICTV taxonomy (*Aquintoviricetes, Polintoviricetes* and *Nucleocytoviricota*-related). This highlights that whilst a useful catch-all term, PLVs are not one viral group. The term largely refers to small eukaryotic DNA viruses which are connected by the capsid morphogenic module (a double jelly-roll fold MCP, a genome packaging A32-like ATPases and an mCP). It also highlights that revisions in these larger viral groups may be necessary to incorporate the PLVs. Using our MCP distance-based clustering as a guide, we focus our analysis on each of the three main groups of PLVs, using the more conserved A32-like ATPase gene to construct detailed phylogenies within each group. Within each of the three groups, ATPase alignments retained considerable phylogenetic signal, with 564–671 parsimony-informative sites per dataset (39.2–54.7% of aligned positions) and 53.1–66.9% of variable sites being parsimony-informative.

**Figure 1.**
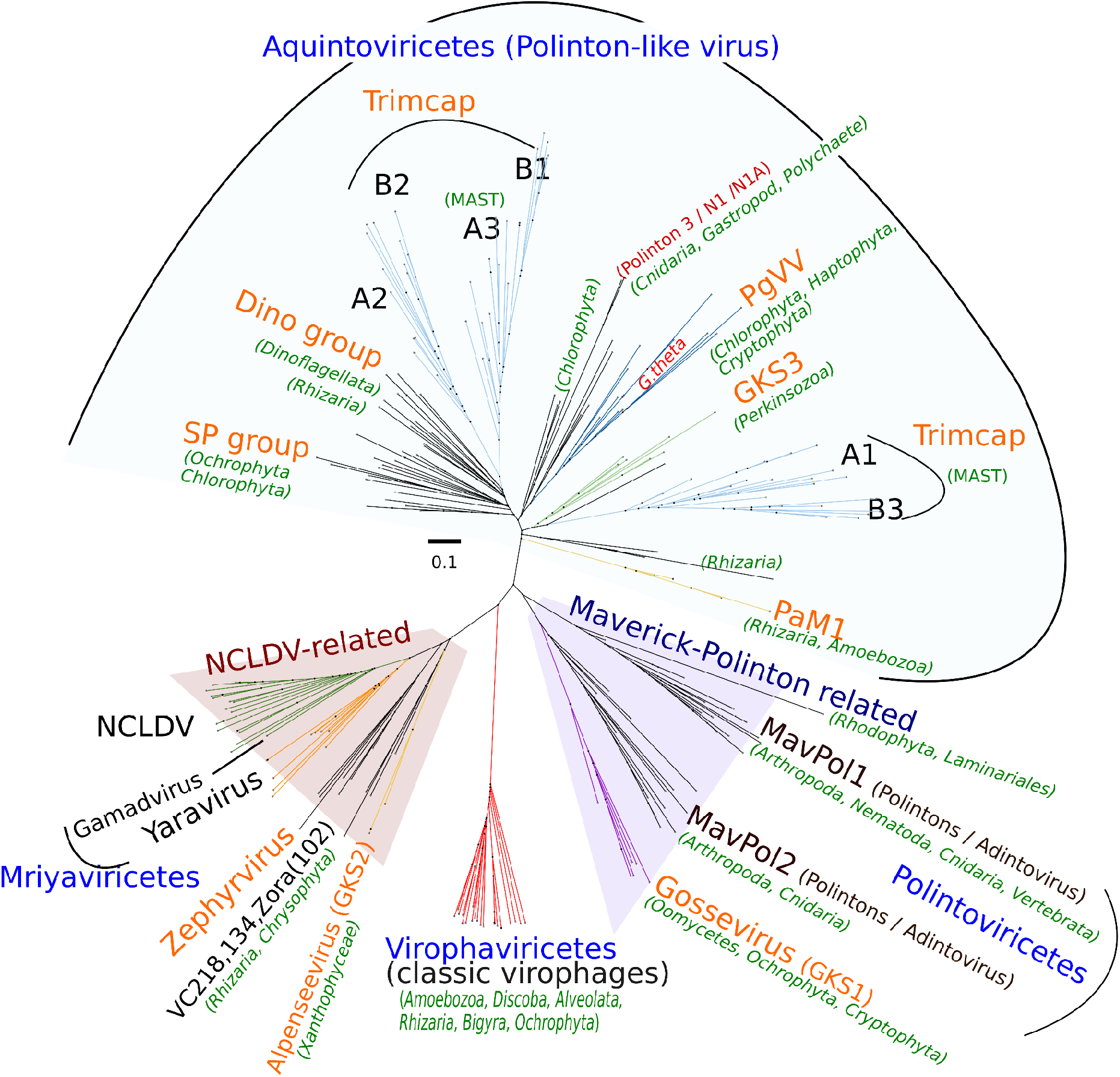
Distance-based clustering of Major Capsid Protein (MCP) HMM profiles showing relationships between Polisuviricots and NCLDV-like virus MCP gene clusters. An all-vs-all HMM profile comparison (HHsearch) of 230 MCP profiles allows for distant relationships to be detected between virophages (Virophaviricetes), Nucleocytoviricota, Yaraviruses, Polintoviruses (Maverick-Polintons) and Polinton-like viruses. HMM profiles were generated from Virus MCP gene clusters with a minimum sequence ID of 30% amino acid identity. Key: NCLDV - Nucleocytoplasmic Large DNA Virus, MavPol - Maverick-Polinton groups. Blue labels are current ICTV viral classes (2025), orange labels are the names of consolidated HMM profiles for new PLV/Mavpol groups provided in the Supplementary Data.

### *Polintoviricetes*-related viruses: Mavericks, Polintons, Adintoviruses and Gosseviruses

Elements described in the literature as Maverick-Polintons fall into the Class *Polintoviricetes* according to the current ICTV taxonomy. Members are almost always found in animal genomes, including many sequences from insects, spiders, molluscs, corals, and vertebrates among others. They have historically been split into two groups [26], which are expanded here by both our MCP clustering (Figure 1) and A32-like ATPase phylogenetic analysis (Figure 2), with both groups found to be associated with similar animal genomes. Group 1 Maverick-Polintons are uniform in terms of gene content, most possess Cysteine proteases (homologous to the adenovirus capsid maturation protease), and pPolB as well as RVE genes, which originally defined Maverick-Polinton elements (Figure 3). Group 2 members have a more variable gene content, particularly in the replication module where pPolB genes are often substituted for superfamily 1 or 3 DNA helicases, which are also generally fused to a DNA polymerase or an AEP family primase–polymerase (PrimPol) (Figure 2 and 3). Our analysis also confirmed that all Adintoviruses clustered within either group 1 or 2 Maverick-Polintons, confirming that the terms Mavericks, Polintons and Adintoviruses are synonymous, which generally refer to animal-centric virus variants as defined by Starrett et al. (2021). Rhodophyte related EVEs are the notable exception to this rule, which were previously found to be related to group 1 Maverick-Polintons [14]. Several other protist groups have been described in the literature as containing Maverick-Polintons, including the oomycete *Phytopthora infestans* and the trichomonad *Trichomonas vaginalis* [2]. This analysis confirms the *P. infestans* ‘Polinton’ as belonging to the Gossevirus group of *Polintoviricetes* (see below). Trichomonad ‘Polintons’, despite our sensitive search strategy, were still unable to be placed in any group owing to a highly divergent morphogenic module, suggesting that further groups will be resolved as more sequences become available.

**Figure 2.**
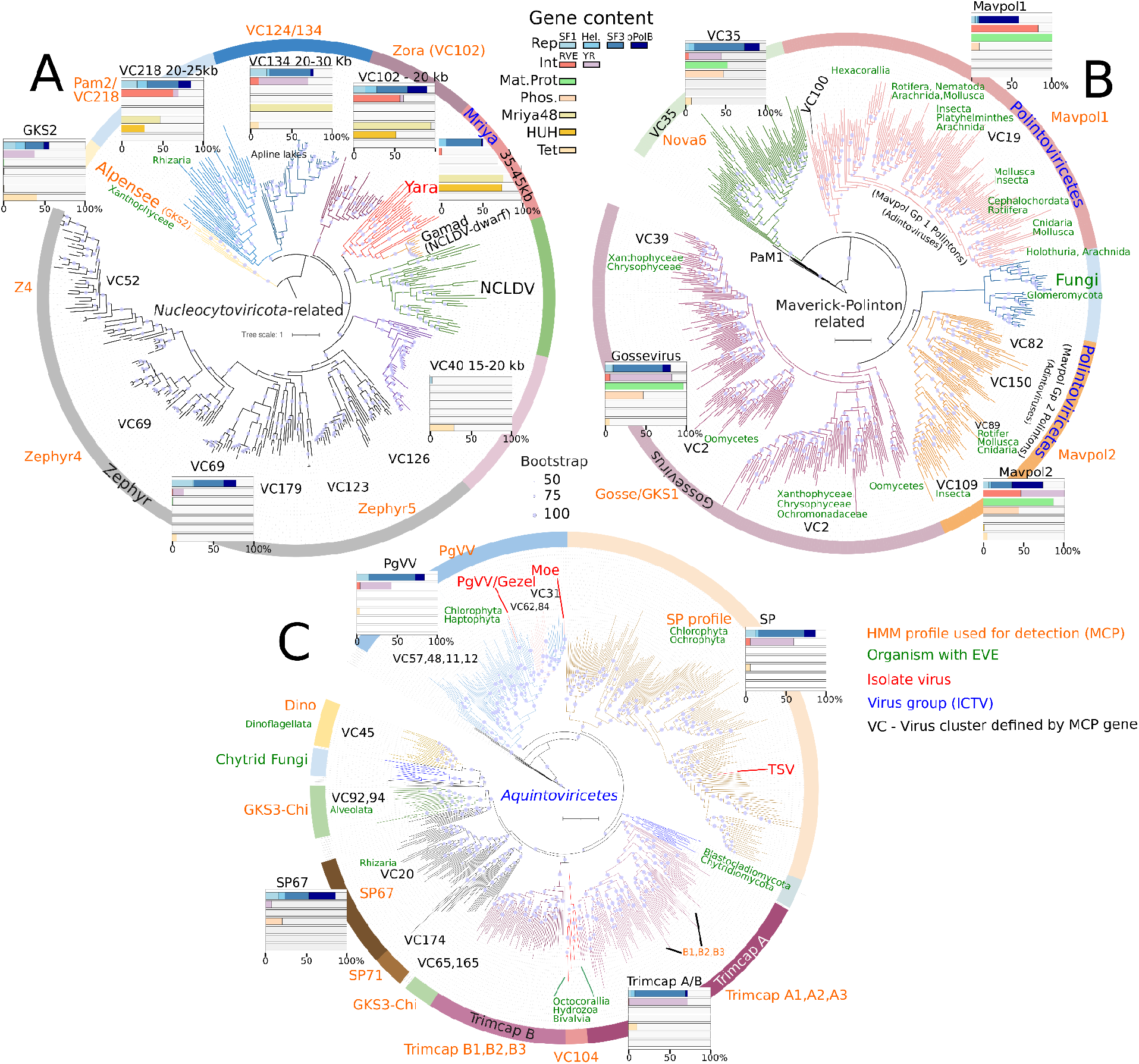
Phylogenetic analysis of A32-like ATPase genes for A) Nucleocytoviricota-related viruses, B) Polintoviricetes-related viruses and C) all other Polinton-like viruses (Aquintoviruses). Virus clusters (VCs) are those defined by our MCP gene clusters (30% amino acid identity), allowing a direct comparison between MCP clustering and ATPase phylogeny. Bar charts for major virus groups represent the percentage of annotated genomes encoding: Rep - Replication gene type (DNA Helicase SF1, unclassified Helicase, SF3 Helicase or pPolB); Int - Integration gene (RVE or Tyrosine Recombinase - YR); Phospholipase (capsid associated gene); Mriya48 gene (Yutin et al 2024); HUH (rolling circle replication gene); Tet - Tet-like demethylase.

**Figure 3.**
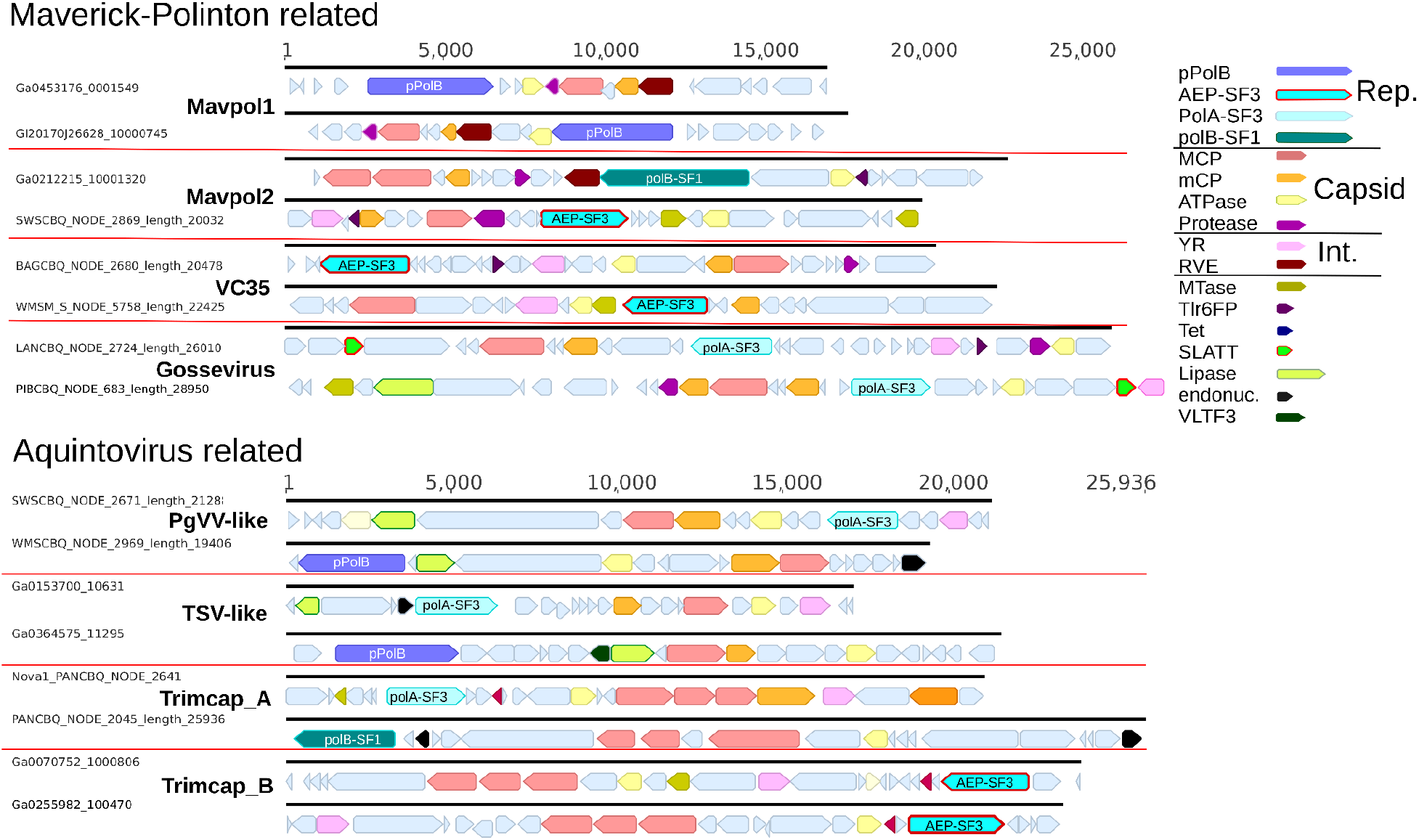
Maverick-Polinton related and Aquintovirus related viruses. Representative genomes are shown from eight groups in Figure 2. Key: pPolB - protein primed DNA polymerase B, SF1-3 Helicase superfamily, AEP - archaeo-eukaryotic primase, MCP - major capsid protein, mCP - minor capsid protein, ATPase - DNA packaging A32-like ATPase, YR - Tyrosine recombinase, RVE - retroviral integrase, MTase - DNA methyltransferase, Tlr6FP - *Tetrahymena thermophila* protein, Tet - Tet-like dioxygenase,, SLATT - SLATT domain, VLTF3 - Virus Late gene Transcription Factor, endonuc - endonuclease.

From our analysis, the Gossevirus group of PLVs was confirmed to be related to the *Polintoviricetes* by both MCP distance-based clustering and A32-like ATPase phylogeny (Figure 1 and 2). This supports a relationship tentatively detected by phylogenetic analysis of a more limited set of MCP genes from alpine lakes [13]. The Gossevirus group is notable in that it is the only major group of PLVs to encode a Cysteine protease as a core gene, further linking them to Maverick-Polintons. Approximately 50% of members also encode a phospholipase domain, which is common in group 2 Maverick-Polintons (Figure 2). Hence MCP, A32-like ATPase and gene content analyses support the conclusion that the Gosseviruses represent a third group of Maverick-Polintons, hence belong within the *Polintoviricetes*. An important distinction, however, is that the Gosseviruses are protist infecting viruses, which are inferred as virus particles in aquatic ecosystems [13], therefore Gosseviruses represent the closest known true virus group to Maverick-Polintons in animal genomes. All Gossevirus MCP genes can be detected by a single HMM profile (GKS1 profile), however, the group is split into at least two main clusters by A32-like ATPase phylogeny (Figure 2), with members from both clusters found throughout Oomycete and Chrysophyte genomes. This suggests they are not specific to a particular host lineage and may be able to infect across different taxa. Gosseviruses are also found in abundance in alpine lakes and other ecosystems, and are actively transcribed by Chrysophytes [13], suggesting they are active viruses in aquatic ecosystems.

To complete the Maverick-Polinton related viruses, a newly identified virus cluster (VC35) was designated based on MCP, A32-like ATPase phylogeny, plus the possession of Cysteine protease and phospholipase genes in approximately half of all members. VC35 members were retrieved from the viromes (<0.2 µm size fraction) of Austrian lakes suggesting they are actively infecting viruses of protists, though the lack of endogenous variants means the host remains unknown. We also identified a number of A32-like ATPase genes from fungal genomes, including *Glomus* spp. and *Rhizophagus* spp., which were phylogenetically related to Maverick-Polinton A32-like ATPases, though these were from fragmented genomic assemblies. Endogenous fungal Maverick-Polintons have previously been detected in *Glomus intraradices* [2], suggesting our findings represent further fungal viruses that group within the *Polintoviricetes*.

### *Nucleocytoviricota*-like viruses: NCLDVs, Mriyaviruses and related PLV

The evolution of giant viruses remains a major question in virology, hence the detection of extant relatives represents an opportunity to further investigate the relationships between PLVs and members of the *Nucleocytoviricota*. Mriyaviruses were recently described as one such group [21], possessing closely related MCP and A32-like ATPase genes to the *Nucleocytoviricota*, plus a number of giant virus orthologous genes including the viral late gene transcription factors 2 and 3 (VLTF2 and VLTF3). This has led to their classification within the *Nucleocytoviricota*, despite their small genome size of 35-45kb [21]. Our analysis of morphogenic module gene phylogenies, plus the possession of a shared set of genes, defines several new groups of smaller DNA viruses related to Mriyaviruses, suggesting further virus groups exist which could be related to the ancestors of the *Nucleocytoviricota*, particularly VC102, which we informally refer to as Zoravirus (from zora, meaning dawn). Members of the Zoravirus group (VC102) and virus clusters VC134, VC218 (Figure 2) possessed genomes in the PLV size range (20-30kb) when considering complete elements (those with direct or terminal inverted repeats). They encoded MCPs, mCPs and DNA packaging ATPases (Figure 4), however, a BLAST search of the Zora A32-like ATPase gene retrieved giant virus ATPase hits from the Genbank non-redundant protein database (ca. 30% identity). Concurrently, Zora representatives pulled from IMG/VR are already catalogued as members of the *Nucleocytoviricota*, despite their small genome sizes, highlighting some shared gene content. Nearly all annotated Zora genomes (20 of 21) encode the core Mriyavirus gene, Mriya48, which also has a homolog in giant viruses as a structural protein in the virion envelope. Approximately half of VC102 genomes also encoded a HUH superfamily endonuclease, a defining characteristic of the Mriyaviruses [21] and essential for initiating replication via the rolling circle mechanism, suggesting many Zora members possess circular replicating genomes. Similarly, VC218 and VC134 members also encode Mryia48 in 47% and 100% of members, respectively, with 28% of VC218 members also encoding HUH endonucleases (Figure 2 and 4). These Mriyavirus-like viral groups differ in their integration genes, with retroviral type integrases (RVE) most commonly encoded in Zora and VC218 members (57% and 62% of genomes respectively), YR most common in VC134 members (60% of genomes), whereas integration genes are notably absent in the Mriyaviruses (Figure 4). The relationships observed by MCP distance-based clustering (Figure 1), A32-like ATPase phylogeny (Figure 2), plus the possession of Mriyavirus core genes across several virus clusters (Figure 4), clearly define several new virus groups which are related to Mriyavirus, hence together, these virus groups represent a link between PLV sized viruses and the *Nucleocytoviricota*.

**Figure 4.**
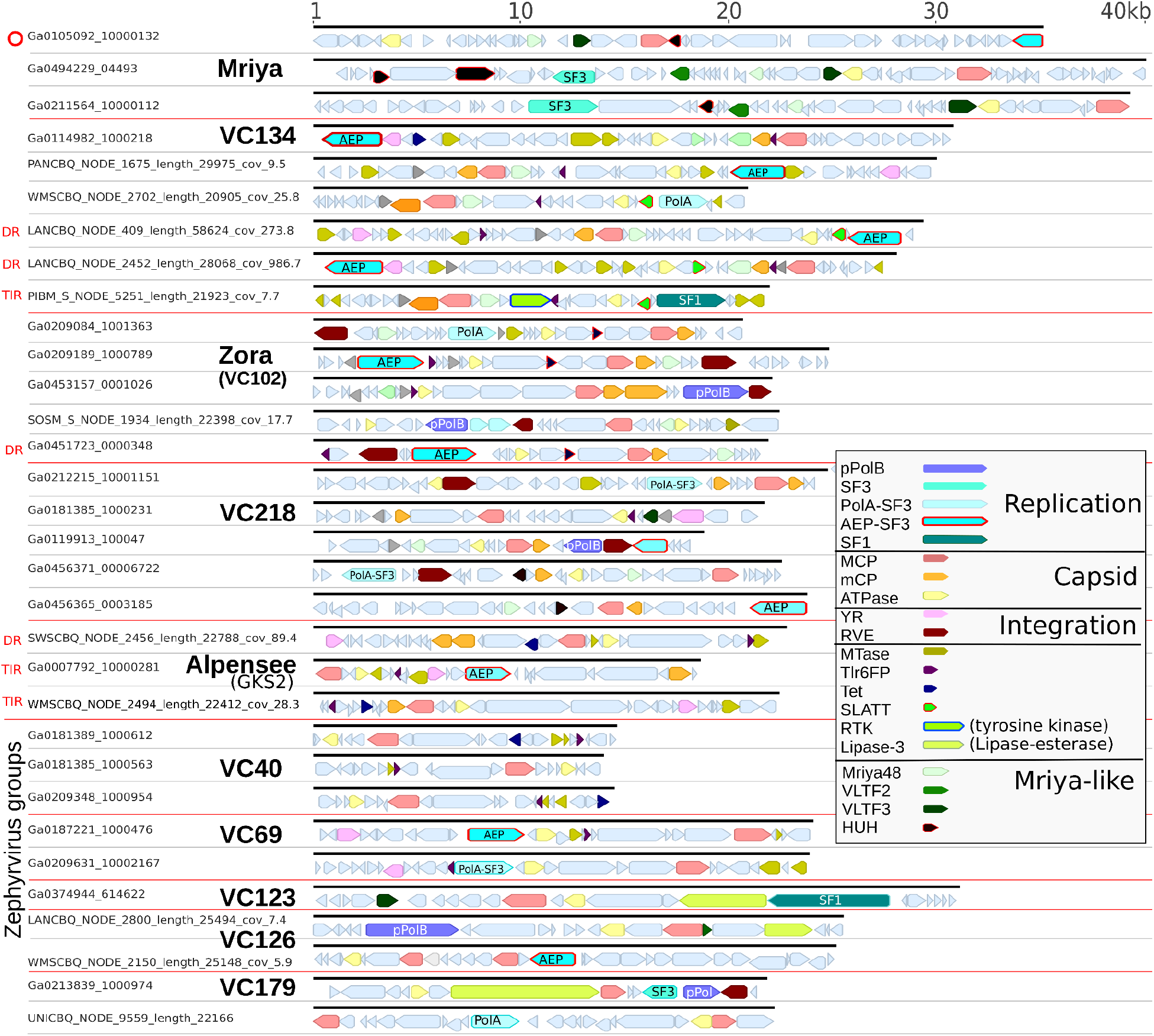
Genome maps of representatives from the NCLDV-like group of PLV, including representative Mriyaviruses. Virus clusters are defined in Figure 2. The circular genome is indicated by a red circle, DR - Direct Repeat, TIR - Terminal Inverted Repeat. Key: pPolB - protein primed DNA polymerase, SF1-3 Helicase superfamily, AEP - archaeo-eukaryotic primase, MCP - major capsid protein, mCP - minor capsid protein, ATPase - DNA packaging A32-like ATPase, YR - Tyrosine recombinase, RVE - retroviral integrase, MTase - DNA methyltransferase, Tlr6FP - *Tetrahymena thermophila* protein, Tet - Tet-like dioxygenase, RTK - Receptor Tyrosine Kinases, SLATT - SLATT domain, Mriya48 - Mryiavirus gene 48 (Yutin et al 2024), VLTF2/3 - Virus Late gene Transcription Factor, HUH - HUH superfamily endonuclease.

Another new and large cluster of PLVs, referred to as the Zephyrvirus cluster (after the west wind of Greek mythology), also share a related capsid morphogenic module to the NCLDVs and Mriyaviruses, but their gene content remains highly variable with few or no Mriyavirus defining genes detected. One subgroup, VC40, (Figure 4) which are also detectable by both Yaravirus-specific and NCLDV specific HMM profiles for the MCP gene (hmmsearch 1e-26 and 3e-25 respectively), contains members with highly reduced genomes of 15-20kb, lacking any detectable replication genes. Only MCP, A32-like ATPase, Tlr6FP and DNA Methyltransferase genes are consistently annotated from this group, with 29% encoding a Tet-like dioxygenase, similar to the GKS2 group of PLV. The main body of Zephyrvirus members (Figure 2) encoded typical PLV replication genes, including DNA Pol A-SF3 fusion proteins - TV-Pol [27], DNA polB-SF1 Helicase fusion proteins, DNA Primase-SF3 Helicase fusion proteins or pPolBs. We also detected VLTF3 genes in approximately 30% of members from the VC123/VC126 clusters, plus a large ORF (up to 7.5kbp) containing a lipase domain in approximately 50% of members (Figure 4). Most Zephyr group genomes were retrieved from freshwater or marine metagenomes in IMG/VR, using two distinct HMM profiles (Zephr4 and Zephy5) to detect the MCP genes, however, several 25Kb elements were also retrieved from the viromes (<0.2 µm size fraction) of Austrian lakes (see VC126, Figure 4), suggesting their existence here as free virus particles. No endogenous examples were found, meaning the hosts for this extensive virus group currently remains unknown.

The GKS2 group, which we refer to as the Alpenseevirus group (Alpine lake virus) to acknowledge their initial detection and abundance in alpine lakes, was also found to be distantly related to Mriyaviruses by the capsid morphogenic module. This distinct group was previously defined by gene sharing networks and MCP phylogeny (Bellas and Sommaruga 2021). Based on updated searches against GenBank WGS (2025), we detected strong MCP hits (up to 72% identity - *Chlorellidium tetrabotrys*) and A32-like ATPase hits (95% amino acid identity - *Chlorellidium tetrabotrys*) to dozens of uncharacterised proteins in the Class Xanthophyceae (Ochrophyta), hence members of the Alpenseevirus group likely infect members of this group and also exist as endogenous viruses (Table S1).

The detection of several new virus groups with capsid morphogenic modules related to Mriyaviruses and NCLDV raises the possibility that these viruses have previously been detected, but binned with members of the NCLDV. To this end, we screened over 58,000 MCP genes from giant virus contigs detected previously from metagenomic data [28], where we detected 518 MCP genes which most closely matched other virus groups (< 1%). These included 254 hits belonging to the recently described Mriyaviruses, 53 belonging to the Zephyrvirus group, 128 to the VC40 group of truncated virus elements and 39 to the Zoravirus group (VC102). Similarly, only 14 of the 1382 MAGs in the GVDB [29] showed evidence of non-giant virus MCP genes, interestingly these included three GVMAGs which included Trimcap group A and B virus members in the bins. Therefore, overall there was only a very low level of PLV or NCLDV-like viruses present in these datasets (Supplementary Table 2 and 3).

### *Aquintoviruses*: Gezel-like, TSV-like and all other PLV groups

All other genomes analyzed fell into previously identified groups of PLV, which are best represented by the current ICTV Class *Aquintoviricetes*. This includes the known isolate, Gezel (PgVV), which exhibits a virophage-like lifestyle [16] and is the type strain of the PgVV group of PLV, which are generally associated with Chlorophytes and Haptophytes. The other published PLV isolate, Tetraselmis striata Virus (TSV) [18] is a member of a much larger PLV cluster, whose members have been detected as endogenous viruses in Chlorophyte and Ochrophyte genomes [14,15]. TSV is a nuclear replicating lytic virus, indicating other members of this group may function as lytic viruses, however, the diverse gene content within all PLV group members hints at a range of life-strategies within each virus group.

The Dinoflagellate associated PLV group (Dino group members), GKS3-Chi and other groups associated with Alveolata and Rhizaria (SP67 HMM profile) aligned within the Aquintoviruses. Interestingly, A32-like ATPase genes from presumed virus regions in numerous Chytrid fungal genomes aligned in two distinct places, supporting previous findings of PLV within the genome of *Spizellomyces punctatus* [13] and suggesting further diverse PLVs are present in these early diverging fungal genomes. Further, some A32-like ATPase genes from Octocorallia, Hydrozoa and Bivalvia genomes were most closely related to Aquintoviruses. These represent the only animal-derived Polintons/PLVs to be found outside the main Maverick-Polinton groups.

The presence of three MCP genes defines the most common type of PLVs in our analysis, the Trimcap (Tri-Major Capsid) group [13]. Members of this group, which each possess three highly divergent MCP genes (Figure 3), were recently recognised as one of the most abundant variants of PLVs in the Southern Ocean [22] (referred to here as group X PLVs). Trimcap viruses were previously found associated with marine Stramenopile (MAST group) hosts based on an analysis of endogenous viral elements [14]. Hence, their true diversity and main hosts are likely found in the marine environment. In this analysis, iterative searches of IMG/VR found that PLVs from the Trimcap group (which we now refer to as Trimcap group A) made up approximately 10% of all retrieved PLV genomes from IMG/VR and approximately 25% of all MCP genes. Group A members could be detected by three specific MCP HMM profiles (Trimcap A1, A2 and A3). However we also detected a second, new group of Trimcap PLVs (Trimcap group B) based on MCP genes found in Austrian lakes. Trimcap group B members possessed MCP genes which were highly divergent from group A, however the A32-like ATPase gene revealed a detectable sequence similarity between the two major groups (Figure 2). Using the A32-like ATPase gene to resolve the Trimcap relationships also overcame the challenge of incorporating multiple MCP genes from each virus genome into a single tree, which were always placed at divergent locations (Figure 1). However, the relationship between MCP and ATPase genes are not always congruent in the Trimcap groups. A subcluster of group A members (as defined by the ATPase gene) contains the group B type MCP genes (Trimcap B1,B2,B4 profiles), highlighting the potential for horizontal gene transfer between the larger PLV clusters. In several Trimcap B members, we also detected two copies of the mCP (penton) gene, meaning up to a third of the PLV genome could encode capsid structural genes. The expansion of the Trimcap groups highlights that multiple MCP genes are a common feature of PLVs and not just a curious oddity. Alphafold 3 modelling of the three Trimcap B MCP genes indicates that whilst each MCP gene can form a homotrimeric assembly, a heterotrimeric capsomer is more structurally plausible (Figure 5), suggesting multiple MCP genes could generate an asymmetric, structurally diverse capsid. The possession of multiple MCP genes is also common across members of the *Nucleocytoviricota* [30].

**Figure 5.**
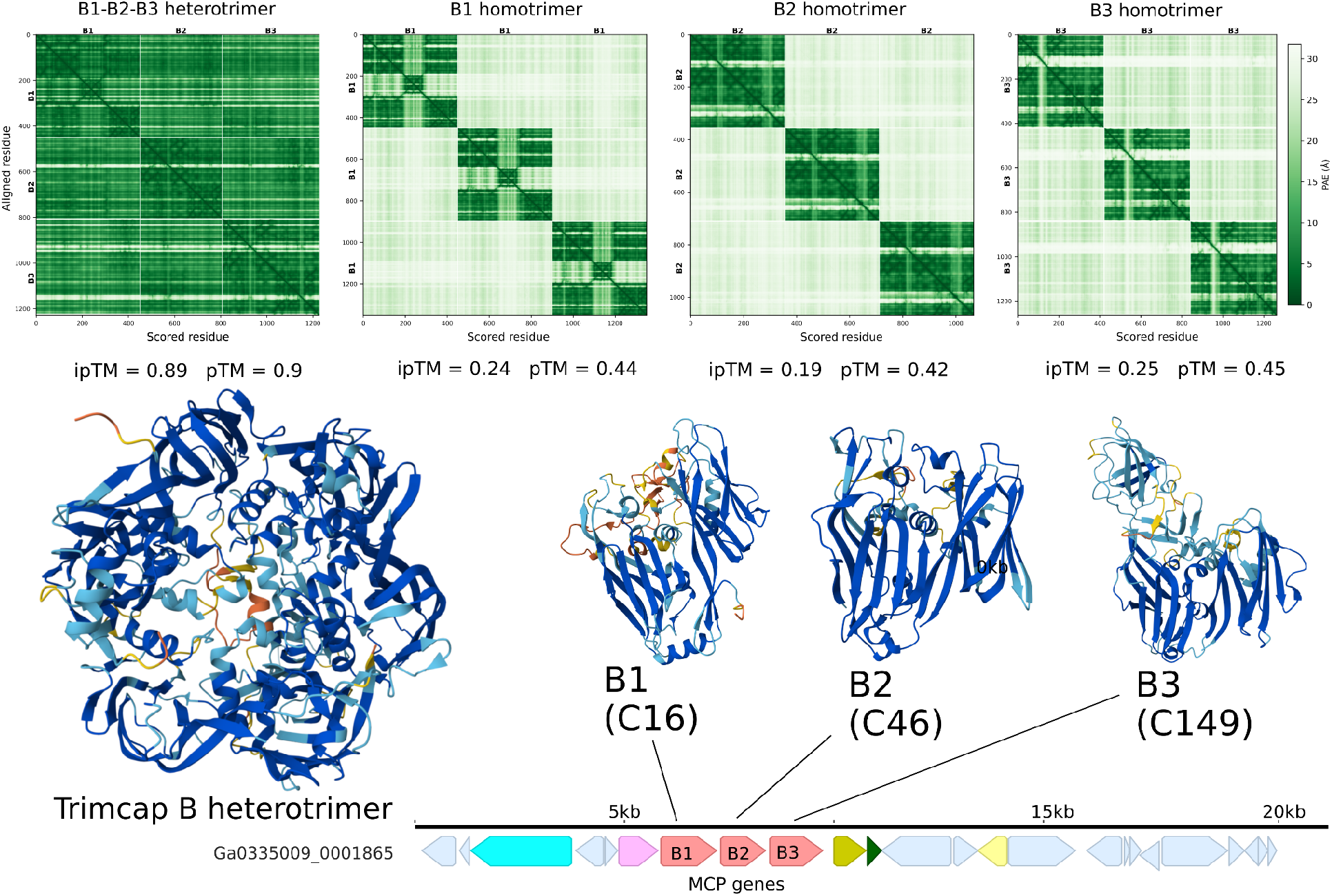
AlphaFold 3 models of a Trimcap group B PLV capsomer. Three highly divergent MCP proteins (B1, B2, B3) were modelled individually and together using AlphaFold 3. The combined model (left) forms a heterotrimeric assembly with high interface confidence (ipTM = 0.89) and low cross-chain Predicted Aligned Error (PAE). In contrast, homotrimeric models of each MCP (models available in Supplementary Data) have weakly supported interfaces (ipTM = 0.19–0.25) and high cross-chain PAE (top). The annotated genome (bottom) is colour-coded as in Figure 4. Key: C16, C46, C149 – MCP types clustered at 30% amino acid identity; B1, B2, B3 – group-level HMM profiles used for detection. ipTM – interface predicted TM-score (confidence in inter-chain interactions); pTM – predicted TM-score (overall model confidence).

#### MELDS and outliers

Mid-sized Eukaryotic Linear DNA viruses (MELDs) as defined by Starrett et al. (2021) were also included in this analysis and found to be scattered across several of the major PLV (Aquintovirus) groups defined here by MCP and A32-like ATPase phylogeny. Hence viruses referred to as MELDs in other studies are included in the large and diverse PLV umbrella.

There were still a number of outlying PLV genomes that were unable to be confidently placed in a particular virus group, despite the increased search sensitivity performed here. These include entomopoxvirus related viruses [25] and Metamonada Maverick-Polintons/PLVs, which remain ungrouped by this analysis. These elements have highly divergent MCP genes which were only confirmed by structural modelling. More virus genomes from related organisms will be needed before they can be linked to the main virus groups.

#### PLV genome content

PLV genomes display considerable genomic flexibility in the replication module, including in elements which encode the same MCP cluster type, further suggesting horizontal gene transfer occurs between many elements. The replication module, which once defined the distinction between Maverick-Polinton and PLV, has been used to infer a direction of evolution between these elements [31]. However, our findings indicate that replication module switches are common across all PLV groups analysed (Figure 2), with multiple different modules present in each virus group. These often switched between a pPolB and an SF1/SF3 helicase which was often fused to a weakly detectable DNA polymerase A or DNA primase domain, including: TV-Pol genes (DNA Pol A-SF3 helicase fusion protein encoded by some virophages) [32]; AEP-SF3 (Archaeo-eukaryotic primase-SF3 Helicase fusion proteins) plus polB-SF1 replication genes. In several virus groups there was no dominant replication module variant, despite members that possessed similar capsid morphogenic modules, for example Mavpol2, Zora(VC102), SP67, VC218 (Figure 2). Such incongruence between the replication module and capsid morphogenic module suggests that horizontal gene transfer must have occurred frequently between PLV groups. The variable replication module within each group of PLV also suggests that different life strategies may be at play within each PLV group; some replication modules may be geared towards giant virus mediated replication, whereas others towards nuclear replication. Previously, MCP phylogeny has been in agreement with whole genome gene-sharing networks created from PLV genomes [13], suggesting the morphogenic module is a more reliable indicator of PLV taxonomy.

Most PLV genomes possessed DNA methyltransferase genes (MTase), with some encoding three or more, which could provide protection from restriction modification systems encoded by coinfecting giant viruses [33]. However, some MTase genes are in proximity to endonucleases, suggesting complete restriction-modification systems which could degrade DNA from coinfecting viruses. Methylation may also be involved in transcriptional regulation. Approximately half of all annotated genomes In the Alpensee (GKS2) group of viruses encoded a Tet-like demethylase, which points towards a role in countering gene silencing mechanisms by the eukaryotic host. Indeed, host methylation of endogenous giant virus genes in the protist *Amoebidium appalachense* has been demonstrated to silence virus gene expression [34], hence methylation could also be important in suppressing endogenous PLVs. The ability to reverse methylation, therefore, may allow PLVs to counter host gene silencing and to enhance their gene expression or reactivation.

## Conclusions

This analysis of PLV relationships, based on the capsid morphogenic module, and combined with the detection of core genes across groups, provides a detailed framework for their future classification. Our work demonstrates that a combination of MCP and A32-like ATPase phylogeny is a useful way to define the Maverick-Polintions, PLVs and Mriyaviruses. Our aim was to untangle the relationships among complex PLV groups, to remove confusion in the literature generated by elements with multiple names and to provide the tools to detect and classify PLV. To this end, we condensed the sequence diversity of PLV and related virus MCP genes into 51 HMM profiles, which can be used for highly sensitive detection of the entire known diversity of PLVs, Virophages, Mriyaviruses and Maverick-Polintons. These profiles will allow users to simultaneously detect and assign any PLV to a particular group based on the highest scoring MCP profile.

## Methods

### Metagenome generation

Viromes were generated from eight Austrian lakes. For each lake, a 20 litre composite water sample was created by sampling from every 1m depth interval. Water was prefiltered though a 142-mm GF/A (Whatman WHA1820150) then a 142-mm 0.22-μm PES filter (Millipore GPWP14250). Viruses were concentrated by iron chloride flocculation as detailed previously [13,35], but without the ethanol precipitation step. Virus DNA was sequenced by Novogene, Cambridge, UK (https://www.novogene.com/). Metagenomic reads were trimmed using Trimmomatic [36] before assembly using SPAdes (settings: --meta -k 21,33,53,77).

**Table 1.** Austrian lake viromes (<0.2µm size fraction) generated and analysed for PLV.

| Lake | GPS | Type | Alt. (m) | Dep. (m) | Prefix | Gb | Contigs (>500bp) | N50 | Assembly (Mbp) | PLV MCP | MCP Mbp<sup>-1</sup> | PLV > 10Kb |
| --- | --- | --- | --- | --- | --- | --- | --- | --- | --- | --- | --- | --- |
| Piburger See 1-12m | 10° 53'19.1" E<br>47° 13'46.0" N | Subalpine mesotrophic | 926 | 24 | PIBCBQ | 55.9 | 97428 | 3781 | 188 | 326 | 1.74 | 109 |
| Piburger See 12-24m | 10° 53'19.1" E<br>47° 13'46.0" N | Anoxic hypolimnion | 926 | 24 | PANCBQ | 68.1 | 320005 | 3032 | 551 | 622 | 1.13 | 239 |
| Reintaler See | 11° 53'36.0" E<br>47° 27'36.0" N | Subalpine mesotrophic | 565 | 9 | RTSCBQ | 55.6 | 340745 | 1645 | 455 | 416 | 0.91 | 47 |
| Schwarzsee Kitzbühl | 12° 22'11.5" E<br>47° 27'27.3" N | Subalpine mesotrophic | 775 | 7 | SWSCBQ | 58.2 | 541774 | 1853 | 772 | 349 | 0.45 | 92 |
| Wildsee Seefeld | 11° 11'29.4" E<br>47° 19'18.5" N | Subalpine mesotrophic | 1177 | 5 | WMSCBQ | 56.1 | 680707 | 1561 | 891 | 593 | 0.67 | 171 |
| Baggersee Rossau | 11° 26'49.0" E<br>47° 15'57.4" N | Lowland eutrophic | 575 | 14 | BAGCBQ | 57 | 488044 | 2013 | 720 | 505 | 0.7 | 122 |
| Lanser See | 11° 25'06.0" E<br>47° 14'25.4" N | Lowland eutrophic | 980 | 12 | LANCBQ | 85.4 | 725306 | 1830 | 1023 | 510 | 0.5 | 162 |
| University pond | 11° 20'28.0" E<br>47° 15'53.0" N | Lowland eutrophic | 600 | 1 | UNICBQ | 69 | 691735 | 4637 | 1390 | 217 | 0.16 | 83 |

### PLV searches, MCP detection and HMM profile construction

To detect new PLV MCP gene clusters, MCP genes from previous studies [13,14] were combined and clustered at 25% amino acid with MMseqs2 [37]. The representative sequence from each cluster with over 10 members (ca. 300) was used to initiate a relaxed iterative jackhmmer search (-N 5, -E 1e-12) of IMG/VR v4 [38] combined with gene predictions from our eight Austrian lake viromes, to create a single file of potential MCP genes. These were then filtered for length (>300aa), then clustered at 30% amino acid identity (mmseqs easy-cluster –c 0.5 –min-seq-id 0.3) generating a representative from each potential MCP cluster. Representatives of known MCP clusters (similar to the input) were quickly retrieved by screening against the initial database using Diamond blastx [39] (settings: –very-sensitive –range culling -F 15, –evalue 1e-12, minimum 23% ID). The remaining clusters (1842) were subjected to a more intensive search to sift out new MCP variants: each representative was used to seed the construction of a refined sequence alignment which could be used to query the pdb70 database. To achieve this, each representative was used to interrogate IMG/VR and the metagenomic database again (jackhmmer -N 1 -E 1e-20). Matches for each representative were filtered (removed sequences >1000 amino acids in length and smaller than the lower quartile of length distributions for the cluster - calculated via seqkit) [40]. Where more than 300 matches remained, they were de-replicated at 70% amino acid identity to reduce numbers below 300 (MMseqs2 easy-cluster -c 0.4). Filtered matches were aligned with MAFFT V7.49 [41] (mafft –maxitertate 1000, –globalpair), with the alignment then converted to a3m format (reformat.py tool in the HH-suite3 package). The a3m format alignment was then used as a hhsearch query (HH-suite3) against the pdb70 database (March 2022 version) (hhsearch settings: -p 20 -Z 250 -glob -z 1 -b 1 -B 250 -ssm 2 -sc 1 -seq 1 -dbstrlen 10000 -realign -mact 0.0 -maxres 32000). A second more sensitive search was then performed on the a3m alignments to emulate an online HHpred search (https://toolkit.tuebingen.mpg.de/tools/hhpred): each of the 1842 a3m alignments was used to initiate a search against UniRef30 (hhblits -e 1e-3 -n 3 -p 20 -Z 250 -z 1 -b 1 -B250) to add in additional, annotated and diverse sequences, before the resulting output alignment was queried against pdb70 using hhsearch as before. This gave two predictions per MCP cluster, MCP hits were confirmed if a positive hit to a viral MCP gene occurred in one or more hhsearch result files, leaving 234 confirmed clusters of PLV MCP genes. To consolidate these clusters further for HMM profile construction, clusters which retrieved more than three overlapping HHsearch hits from the database were merged by combining and re-aligning all members (MAFFT). Alignments were manually inspected, with any poorly aligned sequences removed. This reduced the final number of HMM profiles to 49 which represented Polinton, Virophage, and PLV diversity. For reference, at this clustering level, Virophaviricetes (virophage) MCP genes are represented by a single HMM profile. These profiles were supplemented with five Nucleocytoviricota MCP profiles from the GVOG database (https://faylward.github.io/GVDB/) [29], and two Mriyavirus MCP profiles [21] to allow Mriyaviruses and NCLDVs to be distinguished and filtered from the search hits. These 56 HMM profiles were concatenated and used as our final PLV detection toolkit.

#### Final PLV retrieval

The 56 confirmed HMM profiles for MCP genes were used to perform a full query of IMG/VR v4 and the Austrian lake metagenomes (hmmsearch, bitscore cutoff >75), removing all hits where the top match was to an NCLDV profile. This analysis retrieved 52,658 PLV contigs from IMG/VR, of which 21,145 were over 10kb in length, plus 3598 MCPs in 3328 contigs from Austrian lake viromes (1029 contigs over 10 kb in length). For phylogenetic analysis of core genes and analysis of gene content, these were dereplicated at 97% nucleotide identity across 30% coverage using mmseqs (settings: mmseqs easy-cluster --cluster-mode 3 --cov-mode 2 -c 0.3 --min-seq-id 0.97), leaving 10,937 and 1,025 representative contigs over 10kb from IMG/VR and Austrian lakes which were further analysed.

### Similarity clustering of MCP genes based on pairwise HMM profile comparisons

MCP genes were compared via pairwise HMM profile comparisons to increase the search sensitivity and detect distant relationships. We combined the MCP genes detected above with MCP from previously identified NCLDV and Mriyavirus MCPs [21]. All MCP genes were then built into standardised 30% amino acid identity clusters (MMseqs2) which were aligned using MAAFT as before, creating 234 alignments of MCP genes. Each alignment was built into a HMM model (HH-suite: hhmake) before all files were combined into a custom HH-suite profile database of MCP genes (HH-suite: ffindex_build). Each cluster was then queried against the combined database using hhsearch (all-vs-all hmm profile search) to generate a similarity file. All-vs-all bitscores were converted into a distance matrix using the formula Distance = 1 - (Normalized Score / Maximum Normalised Score), where Normalized Score = Bitscore / Alignment Length and Maximum Normalised Score was calculated based on the highest matching hit pair. A neighbor joining tree was generated from the matrix using Phylip (Figure 1).

### PLV genome annotation

In all retrieved PLV genomes over 10kb, genes were predicted using prodigal (settings: -p meta) before being queried against several databases: Pfam (hmmscan and hhsearch settings: 1e-5), VFAM and VOG databases [42] (hmmsearch 1e-9). Finally, annotated protein clusters from Bellas and Sommaruga 2021 were aligned (MAFFT) and converted into HMM profiles (hmmbuild), these were combined with profiles from Mriyaviruses [21] into a single database. All PLV genes predictions were searched against this final database (hmmsearch -E 1e-9). An annotation was assigned based on a hit to one or more of the databases. Annotations are provided as gf files in the Supplementary Material.

### Phylogenetic alignments of ATPase genes

All genes annotated above as ATPase were extracted from each of the major virus groups defined: Maverick-Polinton-like, Aquintovirus, Virophage and Nucleocytoviricota-like. Sequences were aligned for each group using MAFFT V7.49 (E-INS-I), and a maximum likelihood tree was created using IQ-TREE V2.4 using ModelFinder to select the substitution model (settings:--seqtype AA -nt AUTO -B 1000) [43,44]. The tree was annotated in iTOL (https://itol.embl.de/).

## Data availability

All genomes, alignments and trees used in this study are available at https://doi.org/10.6084/m9.figshare.30112102. Metagenomic data is available in the NCBI BioProject PRJNA1434480. Code and HMM models available to detect PLV at https://github.com/chrisbellas/capscan.git.

## Acknowledgements

We would like to acknowledge Prof. Birgit Sattler and Marie-Sophie Plakolb for assistance with alpine lake sampling. This research was funded in whole by the Austrian Science Fund (FWF), FWF P-34620. For open access purposes, the author has applied a CC BY public copyright license to any author accepted manuscript version arising from this submission

## Notes

https://figshare.com/s/49819b1b91f1ddf41b05

https://github.com/chrisbellas/capscan.git

### Competing Interest Statement

The authors have declared no competing interest.

